# A dual EEG hyperscanning dataset of natural French face-to-face conversation

**DOI:** 10.64898/2026.05.13.724780

**Authors:** Hiroyoshi Yamasaki, Philippe Blache, Daniele Schön

**Affiliations:** Aix Marseille University, CNRS, Laboratoire Parole et Langage (LPL), France; Aix Marseille University, Inserm, INS, Inst Neurosci Syst, France

## Abstract

Conversation is a fundamental human behaviour that requires rapid coordination between speaking, listening, and turn-taking, yet datasets capturing its neural dynamics in natural interaction remain scarce. Hyperscanning EEG is particularly valuable for this purpose because it records both interlocutors simultaneously, enabling the study of speaker–listener coupling, response timing, and dyadic coordination during live exchange. Here we present **DUET** (**D**yadic **U**nderstanding, **E**EG and **T**urn-taking), a hyperscanning dataset for studying natural French face-to-face conversation. The dataset comprises recordings from 18 dyads, or 36 French-speaking adults, performing the Diapix collaborative spot-the-difference task across eight 4-minute face-to-face conversation blocks. For each participant, EEG was recorded from 36 participants; most recordings used 64-channel EEG, with one pilot dyad recorded using 32 electrodes. The public release includes raw EEG recordings, precomputed ICA decompositions for reuse in downstream preprocessing as well as various features derived from the audio and manually corrected transcripts.

## Background & Summary

Conversation is a primary and ecologically natural mode of human language use, in which linguistic information is produced, interpreted, and updated in rapid interaction with another speaker (Levinson and Torreira 2015; Stivers et al. 2009). Rather than unfolding as isolated comprehension or production events, conversation depends on tightly coordinated turn exchange, with interlocutors monitoring one another’s speech and adjusting the timing of their responses. Speaking and listening are therefore deeply interleaved: speakers may begin planning upcoming utterances while still comprehending their partner, and successful interaction depends on the close coordination of these processes (Pickering and Garrod 2013). However, much of the language and cognitive neuroscience literature has examined comprehension and production separately, often in paradigms that reduce or eliminate the reciprocal structure of dialogue (Pickering and Garrod 2014). As a result, the neural basis of spoken interaction in its natural conversational form remains underrepresented in available datasets.

Datasets capable of addressing this gap are rare for both technical and methodological reasons. Recording electroencephalography during overt speech is challenging because articulation introduces muscular and movement-related artefacts, particularly from facial and jaw muscles, which can obscure neural activity of interest (Ganushchak et al. 2011; de Vos et al. 2010). Many EEG studies relevant to dialogue have therefore used third-person or ‘overhearer’ paradigms, in which participants listen to or observe speech exchanges rather than actively engaging in them (Egorova et al. 2014; Egorova et al. 2016; Bögels et al. 2015; Bašnáková et al. 2014; Van Ackeren et al. 2012; Gisladottir et al. 2018). Beyond acquisition, naturalistic conversational data are difficult to analyse because they are continuous, multidimensional, and weakly constrained relative to trial-based tasks. Recent advances in computational approaches, including continuous-time encoding frameworks, have increased the feasibility and value of such datasets, while also highlighting the need for richly annotated, well-structured shared resources (Hamilton and Huth 2020; Zhang et al. 2021; Crosse et al. 2021). A further challenge is that genuine two-person interaction falls within second-person neuroscience, where experimental design and interpretation are more demanding than in single-participant paradigms (Redcay and Schilbach 2019; Hakim et al. 2023). Consequently, only a small number of studies have attempted true EEG hyperscanning during natural spoken conversation (Pérez et al. 2017; Zada et al. 2024), and openly reusable datasets in this space remain scarce.

Here we describe a dual-EEG hyperscanning dataset designed to support the study of natural spoken interaction under conditions that preserve the temporal structure of conversation while remaining suitable for time-resolved neural analysis. Pairs of participants engaged face-to-face in a collaborative Diapix task while EEG was recorded simultaneously from both interlocutors. The dataset includes eight conversational blocks per dyad, together with resting-state recordings, enabling analyses of both interactive speech and non-interactive baseline activity. By combining simultaneous recording from both speakers with a task that elicits spontaneous, goal-directed dialogue, the dataset captures neural activity during turn exchange, overlap, and response preparation in a setting closer to everyday conversation than conventional isolated production or comprehension paradigms.

In addition to the recordings themselves, the dataset provides a derivative layer designed to facilitate reuse across multiple research questions. Speech was transcribed and time-aligned to produce annotation tiers at several levels, including word, syllable, and phoneme timing as well as inter-pausal units (IPUs), linking the conversational stream to discrete linguistic and interactional events. The release also includes acoustic and linguistic features aligned to the neural data, including the speech envelope, fundamental frequency (F0), first and second formants (F1/F2), part-of-speech labels, word surprisal, and word entropy. Together, these derivatives reduce preprocessing and annotation effort and support analyses of both low-level speech structure and higher-level linguistic information within the same shared resource.

This combination of simultaneous dual-participant EEG, face-to-face interaction, and multi-level speech-aligned derivatives makes the dataset suitable for a range of reuse scenarios. The data can be used for encoding analyses relating continuous or event-based speech features to neural activity, for studies of speech planning during listening and turn transition, for investigations of neural processing in more realistic listening conditions than passive laboratory paradigms, and for work on pragmatic and interpersonal coordination during dialogue. More broadly, the dataset is intended as a reusable resource for studying how language processing unfolds when comprehension and production are coupled within interaction, and for supporting methodological development in naturalistic and hyperscanning EEG research.

## Methods

### Participants

Thirty-six healthy adults (mean age = 20.71 years, SD = 3.05; 5 left-handed, 6 males) took part in the study, forming eighteen dyads. Participants were native French speakers with normal or corrected-to-normal vision and no known history of neurological disorders. Within each dyad, participants were unfamiliar with one another prior to the experiment and were matched by sex. Facial hair was used as an exclusion criterion because facial electromyography (EMG) was recorded from electrodes positioned near the mouth. Each participant received 35 Euros for their participation. All procedures were approved by the Inserm Ethics Committee (approval no. 2021-A02248-33). Participants received written study information by email and provided informed consent before participation by email confirmation, in accordance with the Declaration of Helsinki.

### Conversational task and stimuli

To elicit spontaneous conversational exchange, participants completed a collaborative spot-the-difference task based on the Diapix paradigm (Van Engen et al. 2010; Baker and Hazan 2011). In this task, the two members of a dyad view a similar but not identical picture and must identify differences between the images through conversation, without seeing their partner’s version. This paradigm encourages continuous dialogue and frequent turn exchange while providing a shared cooperative goal.

The stimulus set was adapted from the French Diapix materials (Marmel et al. 2020) and supplemented by translated items from the DiapixUK set to complete the full collection. The images depicted three everyday scenes (beach, farm, and street), each represented by four variants, yielding a total of 12 image pairs. Each pair contained 12 differences, although participants were not informed of this number so that they would continue interacting until the end of the trial.

### Experimental procedure

At the beginning of the session, both participants were fitted with 64-channel EEG caps and facial EMG electrodes. The EMG montage consisted of two electrodes placed over the right *orbicularis oris superior*. Participants were then seated face-to-face in the recording room. Electrodes and cables were secured to reduce movement-related artefacts, and participants were instructed to converse naturally while minimising head and body movements as much as possible.

The session began with a 1-minute eyes-open resting-state recording, followed by eight Diapix conversations of 4 minutes each. The eight conversations were randomly selected from the larger pool of 12 image pairs. After the conversational runs, a second 1-minute eyes-open resting-state recording was collected (see Fig. 1). The beginning and end of each section were marked by a short auditory beep. The full session lasted approximately 90 minutes.

**Figure 1.**
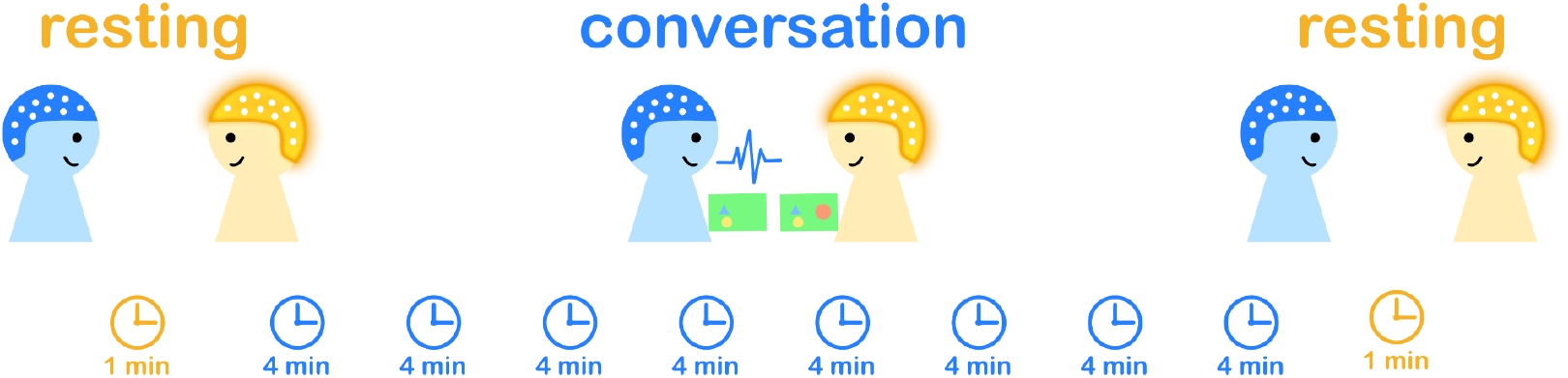
Experimental Procedure. The experiment consisted of eight 4-minute conversations and 1-minute resting-state recordings before and after the conversations.

**Figure 2:**
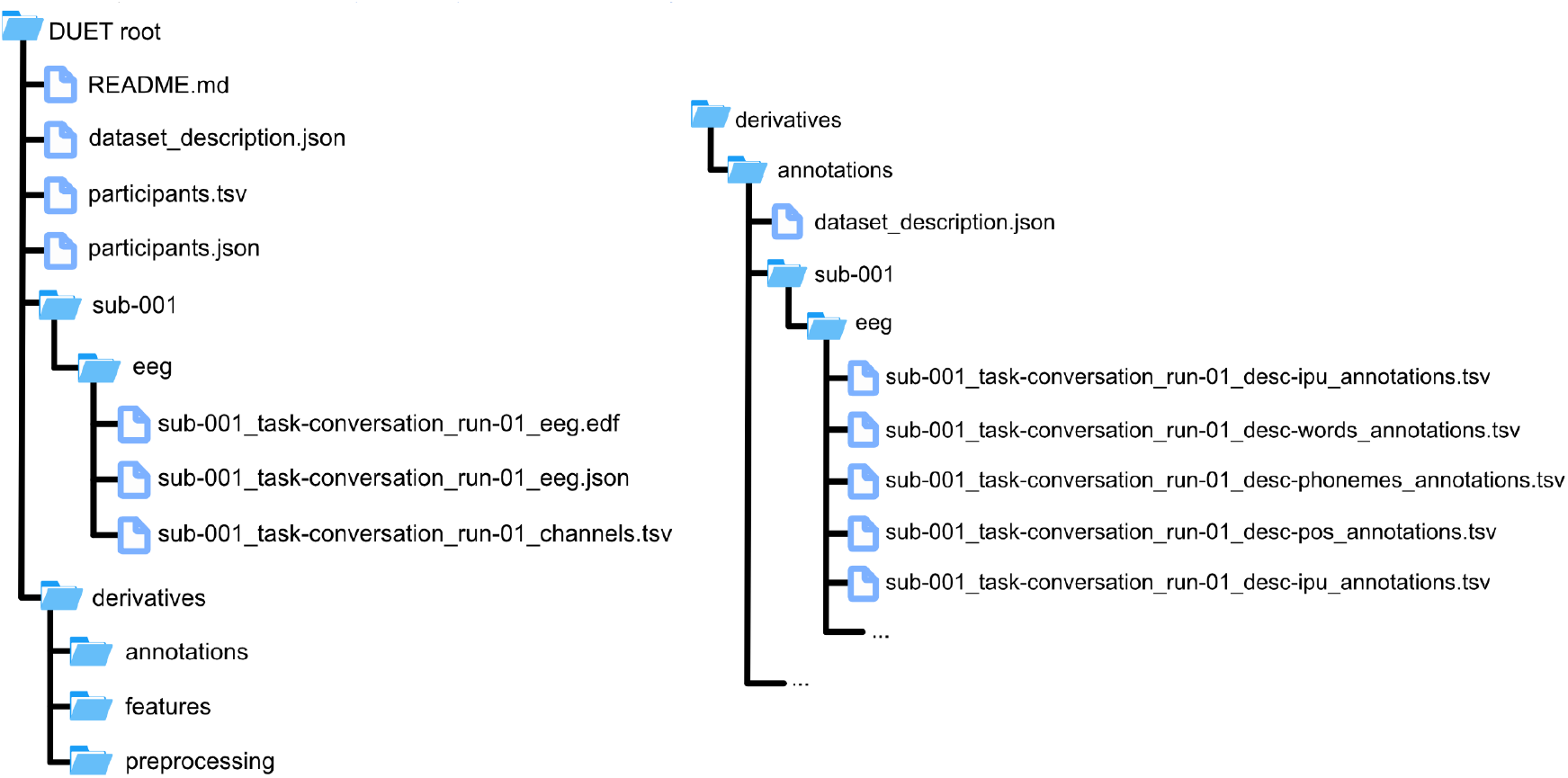
BIDS file structure and annotation directory. Left: Top-level BIDS-compatible directory format. All additional data are contained in the derivatives directory. Right: Annotation directory format.

### Data acquisition

#### EEG acquisition

EEG was recorded simultaneously from both members of each dyad throughout the conversational task using two BioSemi ActiveTwo 64-channel systems arranged according to the standard 10/20 layout (BioSemi, the Netherlands). The two amplifiers were daisy-chained to ensure synchronisation across participants. Signals were DC-coupled, amplified with a low-pass characteristic of approximately -3 dB at 204 Hz, and digitised at 2048 Hz. Earlobe reference channels were also recorded. The trigger signal marking the onset of the beep indicating conversation start, which also serves as the audio-EEG synchronisation anchor (see below), was sent with Presentation® software (Neurobehavioral Systems, Inc., Berkeley, CA).

#### Facial EMG acquisition

Facial EMG was recorded concurrently with EEG using two electrodes placed over the right *orbicularis oris superior*. This recording was included to capture articulatory muscle activity associated with overt speech production during face-to-face interaction.

#### Audio acquisition

Both participants wore AKG C520 head-worn condenser microphones positioned over the EEG caps. Audio was recorded continuously in Audacity (version 3.1.3; Audacity Team 2021) at 48 kHz and 16-bit resolution. Recordings were stored as stereo files, with one speaker assigned to each channel. Audio acquisition was continuous across the session and was cropped manually after the experiment to isolate the relevant task segments.

#### Speech transcription and time alignment

Audio recordings from the conversational runs were cropped to include only the segment from the conversation-start beep through the end of each 4-minute task interval. Transcription was then performed using an automatic transcription pipeline (Yamasaki et al. 2023). The resulting transcripts were manually corrected in Praat (version 6.4.12; Boersma and Weenink 2021). After correction, the speech material was time-aligned at both the word and phoneme levels using SPPAS (Bigi 2015) with the Julius (Lee et al. 2001) backend. The final annotations were stored as Praat TextGrid files for each participant. These annotations provide time-resolved word and phoneme boundaries and serve as the basis for subsequent conversational event definitions and derivative annotations distributed with the dataset.

#### Derived annotations

From the time-aligned annotations, we derived inter-pausal units (IPUs). IPUs were defined as stretches of speech produced by a single speaker and bounded by silences of at least 200 ms. Non-speech vocal events such as laughter and coughs were excluded from IPU boundaries.

Syllable-level annotations were generated from phoneme-level annotations with SPPAS. Syllable annotations contain both exact syllable content (e.g. /s-E/) and syllable-class annotations (e.g. /F-V/, fricative-vowel; for further details see SPPAS documentation).

#### Precomputed ICA models

The provided precomputed Independent Component Analysis (ICA) models were prepared in MNE-Python (Gramfort et al. 2013). To compute the ICA, the data were downsampled to 512 Hz, rereferenced to the average reference, high-pass filtered at 1 Hz, and all eight conversational blocks were concatenated within participant.

ICA was then computed, with the number of components set to the number of channels minus any identified bad channels. Components associated with artefacts were detected automatically using two complementary procedures. First, MNE-Python’s *find_bads_muscle* routine was used to identify muscle-related components based on spectral slope, peripheral topography, and spatial smoothness (Dharmaprani et al. 2016). Second, *ICLabel* was applied as a machine-learning-based classifier of component type (Pion-Tonachini et al. 2019; Li et al. 2022). Components flagged by *either method* were marked as artefactual. On average, 32.1 components were identified for removal (SD = 6.0; see Fig. 3, panel C).

**Figure 3:**
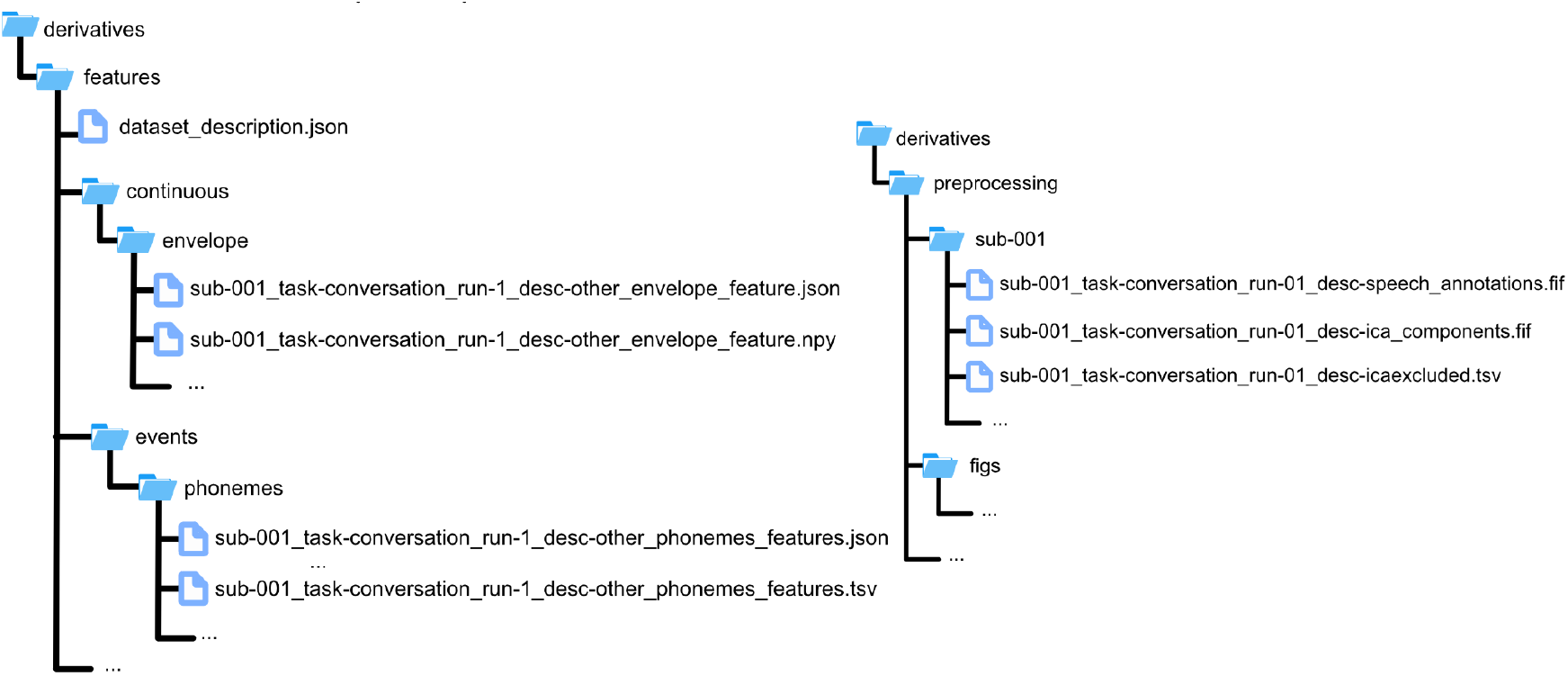
Feature and preprocessing directories. Left: Feature directory format. Features are organised into either continuous or event-like features. Right: preprocessing-related data directory format. It contains precomputed ICA models and MNE-python speech annotations.

For data sharing, the released dataset includes the raw EEG recordings together with precomputed ICA decompositions and component labels. Cleaned EEG time series are not treated as the primary shared data product. Instead, the ICA files are provided as reusable preprocessing provenance, allowing users to inspect, reproduce, or adapt artefact-removal choices for their own analyses.

#### Shared annotations and de-identification

Because conversational speech recordings can contain directly identifiable voice information, the public release does not include the original audio waveform. Instead, the dataset provides de-identified annotation products derived from the recordings. These include manually corrected and time-aligned TextGrid annotations, word- and phoneme-level boundaries, and higher-level conversational annotations derived from these tiers, including IPUs and turn-transition measures.

Identifiable information was removed from shared annotations prior to release. This design preserves the temporal and linguistic structure required for reuse in speech and interaction research while reducing privacy risks associated with public sharing of conversational voice recordings.

### Data Records

The dataset is distributed as a BIDS-organised repository containing participant-level metadata, run-level EEG recordings, and derivative files for conversational annotations, reusable features, and preprocessing provenance. The public release is designed to support reuse for studies of turn-taking, speech planning, dyadic coordination, and time-resolved speech–brain modelling, while avoiding distribution of identifiable waveform audio. In addition to the core EEG data, the repository includes a structured derivative layer containing annotations aligned to the EEG time base and numerical features derived from the transcripts and from privacy-safer acoustic representations. The data are publicly available from OpenNeuro (https://openneuro.org/datasets/ds007764).

### Participant and recording metadata

Participant-level metadata are provided in *participants*.*tsv* (also see Table 1), with variable definitions described in *participants*.*json*. These files contain the anonymised subject information necessary to interpret and reuse the shared dataset. Subject identifiers in the repository correspond to public dataset IDs rather than internal acquisition identifiers. Recording-level metadata are additionally provided through BIDS sidecars and scan tables associated with each participant and run.

**Table 1:** subject list.

| Subject ID | Dyad ID | Age | Sex | Handedness | Runs | Notes |
| --- | --- | --- | --- | --- | --- | --- |
| sub-001 | dyad-001 | 21 | F | L | 8 | 32-electrodes pilot |
| sub-002 | dyad-001 | 19 | F | R | 8 | 32-electrodes pilot |
| sub-003 | dyad-002 | 19 | F | L | 6 | Bad electrode placement, missing first two audio |
| sub-004 | dyad-002 | 18 | F | R | 6 | Missing first two audio |
| sub-005 | dyad-003 | 19 | F | R | 8 |  |
| sub-006 | dyad-003 | 20 | F | R | 8 |  |
| sub-007 | dyad-004 | 26 | F | R | 8 |  |
| sub-008 | dyad-004 | 19 | F | R | 8 |  |
| sub-009 | dyad-005 | 20 | F | L | 8 |  |
| sub-010 | dyad-005 | 19 | F | R | 8 |  |
| sub-011 | dyad-006 | 20 | F | R | 8 |  |
| sub-012 | dyad-006 | 19 | F | R | 8 |  |
| sub-013 | dyad-007 | 23 | F | R | 8 |  |
| sub-014 | dyad-007 | 18 | F | R | 8 |  |
| sub-015 | dyad-008 | 19 | M | R | 8 |  |
| sub-016 | dyad-008 | 27 | M | L | 8 |  |
| sub-017 | dyad-009 | 24 | F | R | 8 |  |
| sub-018 | dyad-009 | 18 | F | R | 8 |  |
| sub-019 | dyad-010 | 32 | F | R | 8 |  |
| sub-020 | dyad-010 | 21 | F | R | 8 |  |
| sub-021 | dyad-011 | 20 | M | R | 8 |  |
| sub-022 | dyad-011 | 18 | M | L | 8 |  |
| sub-023 | dyad-012 | 19 | F | R | 8 |  |
| sub-024 | dyad-012 | 20 | F | R | 8 |  |
| sub-025 | dyad-013 | 19 | F | R | 8 |  |
| sub-026 | dyad-013 | NA | F | R | 8 |  |
| sub-027 | dyad-014 | 20 | F | R | 7 | EEG signal missing in run 8 due to malfunction |
| sub-028 | dyad-014 | 26 | F | R | 7 | EEG signal missing in run 8 due to malfunction |
| sub-029 | dyad-015 | NA | F | R | 8 |  |
| sub-030 | dyad-015 | 19 | F | R | 8 |  |
| sub-031 | dyad-016 | 20 | M | R | 8 |  |
| sub-032 | dyad-016 | 20 | M | R | 8 |  |
| sub-033 | dyad-017 | 21 | F | R | 8 |  |
| sub-034 | dyad-017 | 19 | F | R | 8 |  |
| sub-035 | dyad-018 | 20 | F | R | 8 |  |
| sub-036 | dyad-018 | 22 | F | R | 8 |  |

### Core BIDS EEG dataset

The core release is organised under the standard BIDS subject hierarchy (see Fig. 2, panel A). For each participant, the dataset includes the available raw EEG recordings for the conversation runs, together with the corresponding metadata sidecars and channel tables. These files form the primary electrophysiological record of the experiment. The top-level repository also includes the standard BIDS descriptors required to interpret the dataset structure, including *dataset_description*.*json* and participant metadata files.

### Conversational annotation records

A major component of the release is a derivative annotation layer providing time-aligned descriptions of conversational structure. These annotation files include inter-pausal units (IPUs), phoneme-level alignments, syllable-level alignments, word-level alignments, and part-of-speech tags. All annotation layers are aligned to the EEG time base (EEG conversation-start event), allowing direct integration with electrophysiological analyses and time-resolved encoding models.

Annotations are distributed primarily in tabular form as TSV files accompanied by JSON sidecars that define columns, labels, and controlled vocabularies where applicable. This tabular representation is intended as the canonical public analysis format. These annotations are also provided in Praat TextGrid format for easier visual inspection. The tabular and sidecar structure is intended to make the annotations directly reusable in common analysis environments without requiring custom parsers.

The annotation derivative layer is organised by participant and run within the *derivatives/annotations/* directory (see Fig. 2). Each annotation file corresponds to a specific annotation type and recording run, allowing users to access individual linguistic or conversational layers independently or combine them across levels. For example, users may work only with IPU and turn-transition files for conversational timing analyses, or join word and phoneme records for time-resolved linguistic modelling.

### Derived feature records

In addition to symbolic annotations, the release includes numerical feature derivatives intended for reuse in computational and neurophysiological analyses. These are organised under *derivatives/features/* and divided into feature families according to the type of information represented (see Table 2 for the full feature set).

**Table 2:** Available feature list.

| Feature name | File format | Feature type | Description |
| --- | --- | --- | --- |
| Phoneme | TSV, TextGrid | events | Phonemes generated with SPPAS |
| Syllable | TSV, TextGrid | events | Syllables generated with SPPAS |
| Formants | TSV | events | Median values of F1 and F2 per vowel |
| Word | TSV, TextGrid | events | Word tokens generated with SPPAS |
| Surprisal | TSV | events | Surprisal values computed with Claire Mistral 7B per word token |
| Entropy | TSV | events | Entropy values computed with Claire Mistral 7B per word token |
| POS | TSV | events | Part-of-speech tags added with Stanza |
| IPU | TSV, TextGrid | events | IPU generated from word token annotations |
| Envelope | npz | continuous | Speech envelope array |
| F0 | npz | continuous | F0 values extracted with Parselmouth |

### Acoustic features

The acoustic feature layer contains privacy-safer features derived from speech without distributing the original waveform audio. These include speech envelope, fundamental frequency (F0) and median formants of vowels (F1/F2). These features were extracted with a combination of Praat parselmouth (Jadoul et al. 2018).

The envelope was computed from the speech waveform using a broadband amplitude-envelope procedure (Oganian and Chang 2019). First, the waveform was full-wave rectified and low-pass filtered with a second-order Butterworth filter at 10 Hz using zero-phase forward-backward filtering, yielding a slow-varying amplitude contour intended to capture syllabic-rate modulation. The resulting sample-domain envelope was then segmented into 25 ms analysis frames with a 10 ms frame step, and each frame was summarized by its mean envelope amplitude to produce a frame-aligned contour. No additional frame-level smoothing was applied (smoothing_frames = 1).

Fundamental frequency was extracted from each audio recording after conversion to mono. F0 was estimated using a Praat-based autocorrelation pitch-tracking procedure with a 75–500 Hz search range, 40 ms analysis window, and 10 ms frame step. Unvoiced frames were represented as missing values in the raw F0 contour. For EEG-aligned regressors, unvoiced frames were handled according to the specified fill strategy; by default, unvoiced samples were set to zero. The resulting F0 contour was then aligned to the EEG time axis, using nearest-neighbour alignment when preserving missing values and linear interpolation for filled contours, so that each F0 vector matched the corresponding EEG recording in duration and sample count.

Vowel formants were extracted from pre-existing phoneme alignment intervals. Alignment intervals were read from CSV files, restricted to the target annotation tier, and filtered to retain vowel-labeled intervals. Very short vowel intervals were excluded from estimation using a 30 ms minimum-duration threshold. For each remaining vowel token, formants were estimated from the corresponding audio segment using Burg LPC analysis with a 25 ms window, 10 ms time step, 5500 Hz maximum formant, and up to five formants. F1, F2, and F3 were sampled across the vowel interval, and frames with invalid or missing values were discarded. For each vowel token, the median F1 and F2 were computed across frames with valid F1 and F2 estimates. The output was an event-level table containing one row per vowel interval, including vowel onset, duration, label, speaker, median F1, median F2, and extraction status.

### Linguistic features

Dialogue transcripts were first converted from SPPAS forced-alignment outputs into dyad-level token tables while preserving the original annotation rows and timing information. For each token annotation (*TokensAlign* tier in SPPAS), we retained the original token, speaker label, run number, onset, and offset times, and added fields specifying how the token should be rendered for language-model input. Non-lexical symbols such as @, *, and silence markers (#) were retained in the time-aligned output but excluded from the rendered language-model text, whereas underscore-joined multiword tokens were split into separate lexical pieces for scoring. Subjects were paired into dyads based on subject ID, with odd- and even-numbered participants assigned to speakers A and B, respectively. This procedure produced one token-level CSV per dyad, together with vocabulary and optional transcript-order word files.

For each dyad, we constructed a clean dialogue prompt from the rendered lexical tokens while maintaining a mapping from each rendered character span back to the original SPPAS annotation. Each run was treated as a separate conversational context, and speaker changes, run boundaries, silence markers, and sufficiently long temporal gaps were used to introduce turn or context breaks. Speaker labels were rendered in a bracketed dialogue format, for example *[SpeakerA:]* and *[SpeakerB:]*, following the dialogue-oriented speaker-identification/prefixing conventions used for Claire-style conversational modelling; Claire-Mistral-7B-0.1 is a 7B-parameter causal decoder-only model adapted from Mistral-7B on French conversational data and intended to model dialogue continuations and dialogue-understanding tasks (Louradour et al. 2024). The rendered prompt was then tokenised and scored with a causal language model, yielding position-wise observed surprisal and Shannon entropy. These model-token-level quantities were aggregated back by summing (surprisal) or averaging (entropy) to the rendered lexical word pieces, producing time-aligned informational measures while preserving the original token timings.

Part-of-speech features were generated separately for each participant’s own speech and their conversational partner’s speech. For each run, the aligned token table was filtered by speaker and run, with optional removal of special non-lexical labels before tagging. Token text was reconstructed into a deterministic French text string for POS (part-of-speech) analysis, including normalisation of apostrophes, selected multiword forms, underscores, and punctuation spacing. This text was then processed with a French Stanza pipeline using tokenisation, POS tagging, and lemmatisation (STANZA, Qi et al. 2020). Stanza annotations were mapped back onto the original aligned token rows rather than replacing the original tokenisation. When a token matched a single Stanza token, universal POS tags, language-specific POS tags, morphological features, and lemmas were assigned directly. When one aligned token corresponded to multiple Stanza words, composite tags were stored while preserving the original row. When multiple aligned tokens corresponded to a single Stanza word, POS fields were left empty to avoid arbitrary assignment, and diagnostic mapping labels were recorded. The resulting event-style feature files retained the original token timing and identifiers, added POS, lemma, model, language, and mapping-status fields, and included a JSON sidecar documenting the tagging model, processing settings, and mapping diagnostics.

### Preprocessing and ICA provenance

To support transparent reuse of the EEG data, the repository includes a preprocessing provenance layer under *derivatives/preprocessing/*. This layer contains records related to artefact handling and includes precomputed ICA components and IPU-derived speech segment annotations in MNE-python FIF formats. These files are provided as preprocessing provenance and reusable QC information. They do not replace the raw EEG recordings as the primary electrophysiological data product, and users may choose whether to apply, modify, or ignore the provided ICA information in downstream analyses. They allow reusers to inspect the cleaning history of individual recordings and, where appropriate, reconstruct or modify preprocessing choices.

### File formats and metadata sidecars

Across derivative layers, the release follows a consistent format convention. Event-like and interval-like records are stored as TSV files with JSON sidecars, while continuous sampled predictors are stored in array-based formats with corresponding machine-readable metadata. The sidecars document variable definitions, temporal reference frames, sampling rate where applicable, units, preprocessing steps, model provenance, and any controlled vocabularies used in the annotation or feature records. This format design is intended to make the resource usable across a broad range of analysis environments while preserving enough provenance for reproducibility.

### Release versioning

This Data Descriptor describes version 1.0.0 of the DUET dataset. Version 1.0.0 is the fixed version of record associated with this manuscript and will remain permanently available through the public repository. Any later corrections, additions, or expanded annotations will be released as separate versioned updates and will not replace the version of record described here. Repository metadata and release notes will identify the changes introduced in each subsequent version.

### Data Overview

This section summarises the temporal composition of the recorded conversations to give users a practical sense of what kinds of material are available in the dataset. In natural dialogue, participants alternate between self-produced speech, listening, silence, and occasional overlapping speech, and these interactional states are not equally suitable for all downstream analyses. This is particularly important for EEG, where overt speech is associated with substantial articulation-related artefacts (de Vos et al. 2010), making analyses more reliable during listening or non-speaking intervals than during self-produced speech. For this reason, it is useful to characterise not only the annotation structure itself, but also the relative amount and temporal organisation of different conversational states represented in the corpus.

To this end, we summarise several descriptive properties of the conversations (see Fig. 4). The distribution of inter-pausal unit (IPU) durations characterises the range and variability of contiguous speaking segments and indicates the temporal scale of production events available for analysis. Cumulative speaking-time measures show how vocal contribution accrues for each participant over the course of an interaction, providing a descriptive view of relative participation within dyads. In addition, the breakdown of speech, silence, and overlapping speech quantifies the temporal composition of the recordings and helps indicate how much material is available for analyses focusing on speech production, listening periods, turn-taking, or cleaner non-speech intervals. Together, these summaries document the artefact structure of the recordings and the behaviour of the provided ICA decompositions, allowing users to evaluate whether and how to apply the shared ICA information in their own preprocessing pipelines.

**Figure 4.**
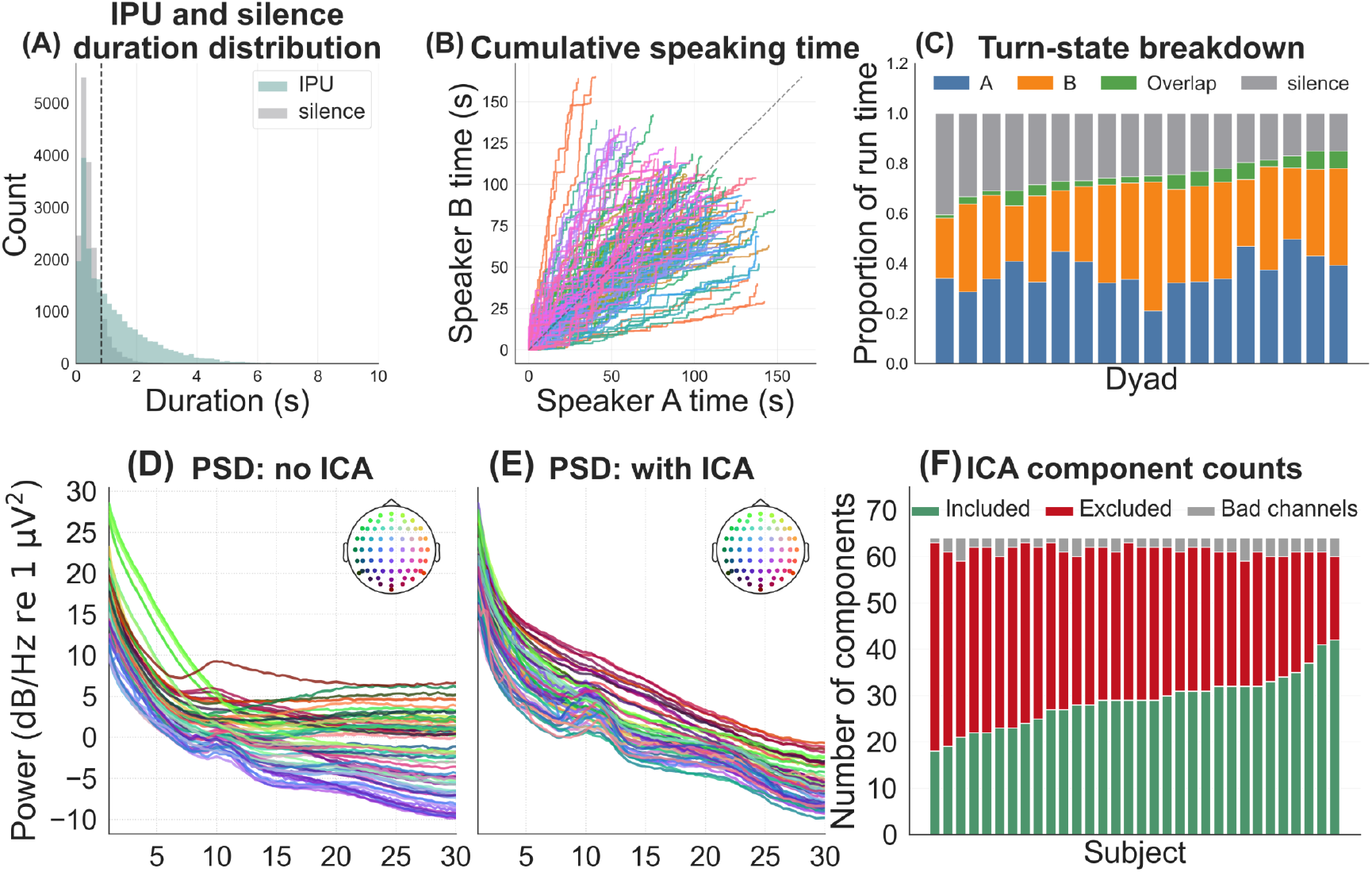
Dataset description. (A) distribution of inter-pausal unit (IPU) durations pooled across dyads and runs, with the dashed line marking the median duration. (B) cumulative speaking time for speaker A versus speaker B within each run; the dashed diagonal indicates equal cumulative speaking time, so traces above or below the line reflect asymmetries in participation. (C) per-dyad proportions of run time spent in four turn states, showing periods where only speaker A spoke, only speaker B spoke, both speakers overlapped, or neither speaker was speaking. (D) grand-average power spectral density before ICA, showing the overall spectral profile across channels. (E) grand-average power spectral density after ICA, allowing comparison of spectral attenuation following artefact removal. (F) per-subject decomposition of ICA outcomes, showing the number of retained components, rejected components, and bad channels, which summarises how much cleaning was required for each participant.

### Technical Validation

Technical validation was designed to assess the usability and internal consistency of the released data and derivatives, rather than to test hypotheses about conversational language processing. We evaluated two aspects of the release: first, whether the EEG recordings and accompanying ICA provenance support plausible artefact attenuation; and second, whether the released speech-aligned predictors are temporally aligned with the EEG well enough to support standard continuous encoding workflows. To this end, we report descriptive summaries of ICA cleaning and benchmark temporal response function (TRF) analyses as quality-control checks (Crosse et al. 2021).

Because overt speech generates substantial movement, muscle, and articulation-related contamination in scalp EEG, conversational datasets require explicit validation of the cleaning procedure. We therefore summarised the effect of independent component analysis (ICA) cleaning in two ways. First, we quantified the number of ICA components retained and excluded across recordings, providing a transparent overview of how strongly the data were affected by stereotypical artefacts (see Fig. 4 panel F). Second, we compared power spectral density (PSD) estimates before and after ICA cleaning. These spectra were used as a sanity check that preprocessing attenuated artefact-related power (see Fig. 4 panels D and E), particularly in frequency ranges typically associated with muscle contamination, while preserving the overall spectral structure expected of physiological EEG. Together, these summaries are intended to show that the the released raw EEG and ICA provenance support a plausible preprocessing workflow for downstream analyses

The data used in the following analyses were downsampled to 64 Hz, rereferenced to the common average, cleaned by applying the precomputed ICA described above, interpolated for manually annotated bad channels, and band-pass filtered between 1 and 30 Hz.

We next used forward TRF encoding models as a benchmark quality-control analysis to assess whether the released EEG recordings, timing information, and speech-derived predictors preserve recoverable stimulus-aligned structure after applying one documented preprocessing workflow. For each participant and conversation run, preprocessed EEG was cropped to 4 minutes each. Speech-derived predictors were aligned to the same temporal grid. Continuous features, such as the interlocutor’s speech envelope and F0, were resampled directly, whereas event-like linguistic features, including phoneme, syllable, token, function-word, content-word, surprisal, and entropy onsets, were converted into sample-space impulse-based regressors. Composite predictor families were expanded into their constituent design columns before model fitting.

TRFs were estimated using a forward encoding model that predicted multichannel EEG from time-lagged speech features over a lag window from -200 ms to 1000 ms using sPyEEG (Guilleminot et al. 2026). To avoid temporal leakage across adjacent samples, each cropped run was divided into four contiguous blocks, and lagged design matrices were constructed separately within block boundaries. Model fitting used nested grouped cross-validation with ridge regression as implemented in sPyEEG’s *TRFEstimator*. Outer folds were grouped by run and inner folds by segment. For each outer fold, the regularisation parameter was selected from a 50-point log-spaced grid from 10^−2^ to 10^8^ based on inner-fold performance, after which the model was refit on the outer-training data and evaluated on held-out outer-test data. Predictive performance was quantified as the mean channel-wise Pearson correlation between observed and predicted EEG and then averaged across outer folds to obtain a subject-level score.

To determine whether model performance reflected genuine stimulus–response coupling rather than trivial temporal structure, we compared real-model performance with a null model based on circularly shifted envelope predictors. Shifts were applied independently within each run after temporal alignment, preserving the autocorrelation structure of the predictors while disrupting their true correspondence with the EEG. Above-null performance in this benchmark was interpreted as a quality-control indication that the acquisition, synchronisation, preprocessing workflow, and predictor alignment are mutually consistent enough to support continuous encoding analyses. This comparison was not used to test a specific cognitive or linguistic hypothesis.

Finally, we assessed the usability of the released feature set through feature-ablation analyses. A full model including all configured predictor families was refit repeatedly with one predictor family or feature group removed at a time. For each comparison, we computed the subject-level ablation delta as the full-model held-out prediction accuracy minus the score obtained after removing that feature group. These deltas quantify the extent to which each derivative contributed non-redundant predictive information within the documented TRF workflow. To assess whether these contributions were reliable across participants, subject-level deltas were tested against zero using two-sided one-sample t-tests. This analysis was used as a quality-control procedure for the released derivatives, rather than as a hypothesis test about the cognitive or linguistic importance of individual feature classes. As shown in Fig. 5B, all tested feature groups except word onset showed ablation deltas significantly greater than zero, indicating that they contributed non-redundant predictive information within this benchmark model. In addition, TRF coefficients were aggregated across folds and subjects to generate kernel visualisations for selected predictors (Fig. 5C). These kernels provide an additional sanity check by showing whether fitted weights exhibit stable, temporally structured profiles rather than unstable or noise-like estimates under the benchmark model.

**Figure 5.**
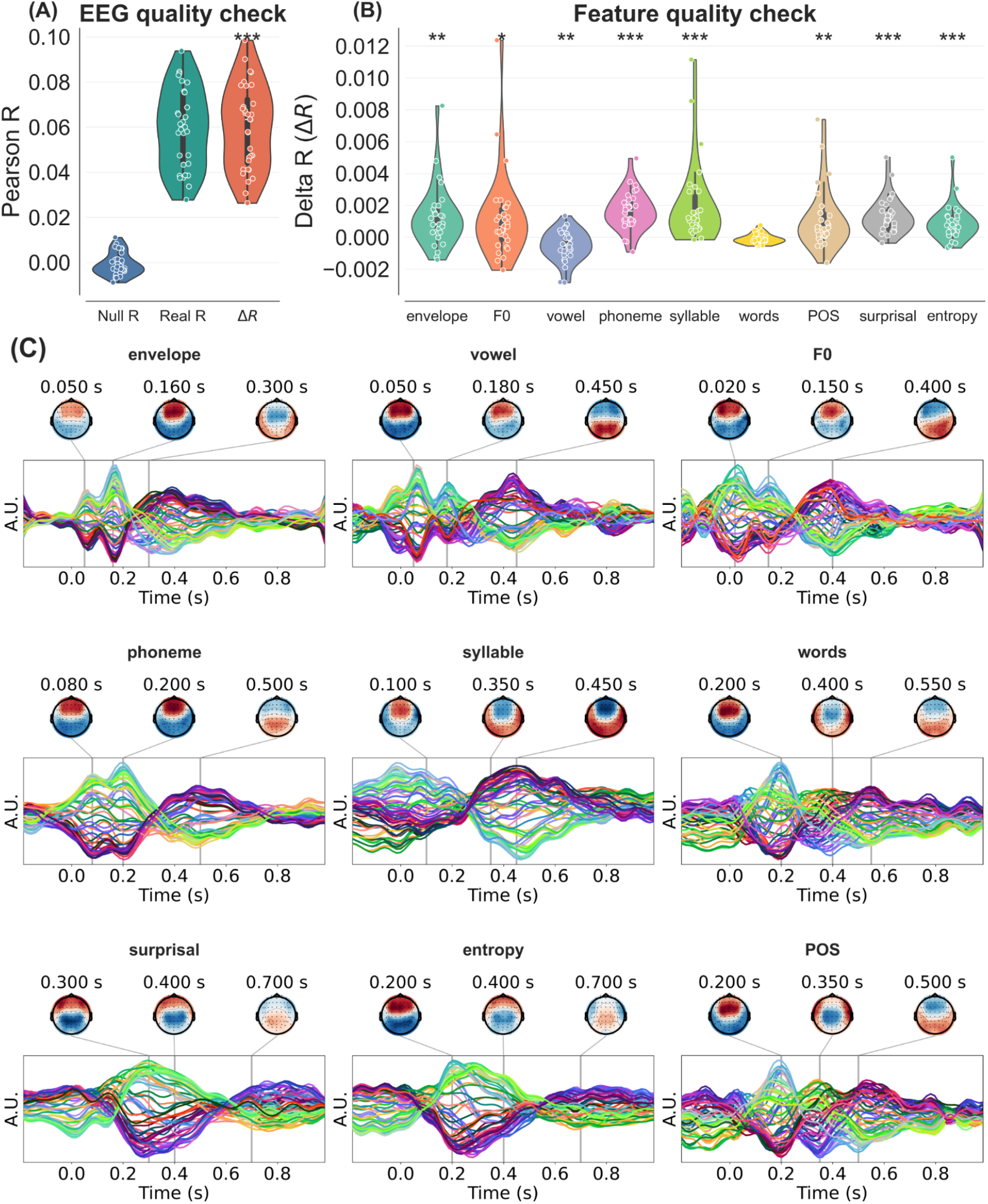
TRF model quality-control summary. (A) subject-level comparison of held-out prediction accuracy for the real model, the null model, and their difference, showing whether the fitted TRF captures reliable EEG–speech relationships beyond chance. (B) feature-ablation results, where each distribution shows the drop in held-out performance when one predictor is removed from the full model; larger positive values indicate greater unique contribution of that feature to model performance across subjects. (C) the temporal response function for each predictors, with the line plot summarising response magnitude over lag and the scalp maps showing the corresponding spatial topographies at selected time points. Together, the panels illustrate how different acoustic and linguistic features evoke distinct temporal and spatial response patterns in the EEG.

Taken together, these preprocessing summaries and TRF-based benchmarks support the technical validity of the dataset at two levels. The preprocessing diagnostics indicate that conversational EEG artefacts can be attenuated while preserving plausible electrophysiological structure. The encoding analyses provide a benchmark demonstration that, after applying the documented preprocessing workflow, the released EEG, timing metadata, and derivative predictors can be combined in a standard continuous-modelling pipeline. These results are presented as technical validation of data usability and alignment, not as evidence for a particular model of conversational language processing.

### Usage Notes

This dataset is intended to support reuse in studies of naturalistic spoken interaction, particularly for analyses of turn-taking, speech–brain coupling, and the temporal organisation of listening and speaking during conversation. Because the release includes time-aligned conversational annotations and derived acoustic and linguistic features, it can also be used for event-related analyses, temporal response function (TRF) modelling, and representational analyses based on word-level surprisal, entropy and part-of-speech labels.

Raw waveform audio is not distributed with the public release because participants’ voices may be identifying. The public dataset instead provides de-identified, time-aligned annotations and derived feature files as the reusable representation of the speech stream. Users should therefore treat the annotation and derivative layers as the primary speech-related data products. The dataset is well suited to analyses of conversational timing, turn exchange, listening periods, and speech-aligned neural responses, but it is not intended for analyses that require direct access to the original waveform or speaker-identifying acoustic properties.

Users should also take into account that several shared linguistic derivatives are model-dependent rather than direct observables. In particular, surprisal and entropy depend on the exact language model, tokeniser, revision, context window, and aggregation procedure used during feature extraction. These quantities should therefore be interpreted as reproducible outputs of a specified computational pipeline, not as model-independent properties of the stimulus. For this reason, each derivative should be used together with its accompanying metadata, including model and tokeniser revision, preprocessing rules, context length, layer specification where relevant, sampling rate, units, and aggregation rules recorded in JSON sidecars.

For EEG analyses, users should consider residual speech-production contamination when analysing periods in which participants are overtly speaking. The dataset is especially well suited to analyses that explicitly distinguish between speaking, listening, silence, overlap, and listening-only windows, and these distinctions should be incorporated into analysis design where relevant. For example, researchers interested in minimising articulation-related artefacts may preferentially focus on listening-only segments, whereas researchers studying dialogue dynamics may use the turn-structure annotations to model gaps, overlaps, and response timing directly.

More broadly, the release was designed to remain maximally reusable despite the absence of shared audio. In practice, this means that users can still perform a wide range of modern speech neuroscience analyses, including encoding and decoding models, analyses of conversational structure, and tests of linguistic prediction using transcript-derived features. The dataset and derivative structure were also chosen to support low-friction reuse through permissive licensing and explicit versioning of computational feature pipelines.

## Code availability

Code used to compute the features and ICA is available at:

Main code: https://github.com/CogSci-hiro/duet

VoxAtlax: https://github.com/CogSci-hiro/voxatlas

Data are available at:

https://openneuro.org/datasets/ds007764

## Acknowledgements

We thank the participants for their commitment, our team members for their help with transcription correction, Chiara Mazzocconi for help with annotations, Sophie Chen for help in the experimental setup and data acquisition, Emma Berthault and Anne Mathieu for help with data acquisition. Research was supported by grants ANR-21-CE28-0010 (DS), ANR16-CONV-0002 (ILCB), ANR-17-EURE-0029 (NeuroMarseille), and the Excellence Initiative of Aix-Marseille University (A*MIDEX).

## Author contributions

Conceptualisation: DS, PB Data acquisition: HY

Data curation: HY Formal analysis: HY

Funding acquisition: PB, DS Project administration: DS Visualisation: HY, DS

Writing – original draft: HY, DS

Writing – review and editing: HY, PB, DS

